# Isolation and Transfection of Rice Egg Cells and Zygotes for Cellular Localization

**DOI:** 10.1101/2022.08.19.504568

**Authors:** Hengping Xu, Laura Bartley, Marc Libault, Venkatesan Sundaresan, Hong Fu, Scott Russell

## Abstract

Due to the difficulty in accessing to gametes and zygotes in flowering plants, which are controlled in single cells deeply embedded in multiple tissues, little is known about how the initiation of plant embryogenesis may reflect or contrast from such systems in other eukaryotes. In this study we has developed an approach of isolation and transfection of rice egg cells and zygotes for cellular localization of rice cell cycle factors (KRP5, KRP4 and FB3), which opened a pathway to monitor protein expression in rice egg cells and zygotes at different developmental stages. The advantageous feature of isolated rice cells may serve as an ideal system for studying the molecular mechanism underlying the rice zygotic division to initiate seed formation.

## Introduction

The plant sexual reproduction is one of fundamental issues for human. It includes the double fertilization as the central stage in the life cycle of higher plants, referring one sperm cell binds and fuses with the egg cell then develop into the embryo, and the other with the central cell into the endosperm for the nutrition of the embryo. Both embryo and endosperm are the two major components of seeds to feed human and animals. This unique process was firstly described in 1898 by Sergius Nawaschin and 1899 by Leon Guignard in Lilium martagon (Hu 1998; Russell 1992). Since then, intensive research has been devoted to the observation and characterization of the diverse reproduction process and the method invention particularly in electron microscopy to rich the knowledge of gametogenesis and embryogenesis in angiosperms. However, the molecular mechanisms underlying the double fertilization, the gamete formation and zygote development have been very limited due to inaccessibility to these cells deeply embedded in multiple plant tissues as well the lower expression level of those major biochemical components in the process for the seed formation.

Therefore, the great effort with significant progress has been made in the manual isolation of plant gametes, mainly sperm cells, egg cells, and zygotes during the past several decades (Lin et al. 2020; Russell 1985; Russell et al. 1990; Theunis 1991; Zhang et al. 2010).

One of the earliest attempts in isolating plant sperm cells can be tracked back to 1973 (Cass) from barley, *Hordeum vulgare*. The spherical shape was observed after released from pollen grain and used for observing the ultrastructure. However, the sperm cell isolation in mass has not been achieved until 1986 from *Plumbago zeylanica* (Russell 1986). Since then, the sperm isolation has been undertaken using at least 29 different plant species, including those with economic importance such as maize (*Zey mays*), rice (*Oryza sativa*), wheat (*Triticum aestivum*), rape (*Brassica spp*), spinach (*Spinacia oleracea*), lily (*Lilium longiflorum*) and leek (*Allium tuberosum Roxb*). In general, the sperm cells were released from pollen grains or pollen tubes broken by grading or osmotic shocking, then enriched using Percoll gradient centrifugation. The isolated sperm cells had been used for identifying sperm specific proteins (Blomstedt et al. 1992; Southworth and Kwiatkowski 1996; Xu and Tsao 1997), sperm specific transcripts (Xu H et al. 2002; Xu HL et al. 1999) and *in vitro* fertilization (Faure et al. 1994; Kranz and Lort 1994).

The viable embryo sacs and egg cells were isolated during 1980s in China from several plant species including *Antirrhinu majus* (Hu et al. 1985). Then, a large quantity of female gametes was isolated in Europe and America (Cao and Russell 1997; Huang and Russell 1989; Huang et al. 1990; Kranz et al. 1991; Mol 1986; Van and Kwee 1990; Wagner et al. 1988). The number of plant species used for this purpose are over 48, including 12 crop plants. The major isolation technique was manual micro-dissection with or without the aid of cell wall enzymatic digestion. The micro-dissection needs micro-knives, thin needles, micro-capillaries and inverted microscope. Without enzymatic digestion in this method, it is easier to control the quality and purity of isolated products, but the disadvantage is that it requires great skill and time consuming. With aid of enzymatic digestion in a mixture of cellulase and pectinase to soften the surrounding tissues, a large quantities of female gametes can be isolated, the potential issue is hard to completely to avoid residual degrading effect on the viable cells (Dumas and Mogensen 1993; Hu et al. 1985; Kranz et al. 1991).

As one of staple crop plants, the rice has great potential in helping us meet the challenge of increasing global population. Meanwhile, it is the best model of monocot plants with sequenced genome and excellently documented record in agronomy, genetics and cell biology. That is why it is important to develop the procedures to isolate rice gametes and zygotes to facilitate the study of the mechanism underlying the rice seed formation for its better production in the future.

To our knowledge, the first mass isolation of rice sperm cells is reported in 1999 (Gou et al 1999; Russell et al 2017). In the same year, the method was developed to isolate rice egg cells and zygotes and the isolated zygotes were used for *in vitro* culture to generate fertile rice plants. (Zhang et al. 1999; Zhao et al. 2000). In addition, the similar dissection but with more careful manual manipulation was applied in isolating the larger central cells (Uchiumi et al. 2006; Zhao et al. 2000).

The isolated rice gametes were used for *in vitro* fusion for artificial zygote (Khalequzzaman and Haq 2005), and detection of specific gene expression (Ohnishi et al. 2011). The isolation of rice gametes and zygotes were also systematically optimized for higher purity and quantity in the Russell Lab and the products were applied for a number of studies, including *cis*-regulatory elements and transcriptomes of rice sperm cells and egg cells (Anderson et al. 2013; Russell et al. 2012; Sharma et al 2011), large-scale transcriptomic changes in rice zygotes (Anderson et al 2017) and the landscape of small interference RNA (siRNA) in rice gametes and zygotes (Li et al. 2020 and 2022).

However, it is rarely reported about using the isolated rice gametes and zygotes in the transfection for cellular localization. Here we present the result of isolation and transfection of rice egg cells and zygotes for cellular localization of three rice cell cycle factors (KRP5, KRP4 and FB3). The feature and potential application of this procedure for molecular study in plant reproduction biology will be discussed.

## Material and Methods

### Plant material

The seeds of rice (the variety *Kitaake*) are treated with 20% bleach for 10 min and followed by 3 washes with autoclaved water, 10-15 minutes each, then germinated with sterile water in petri dishes in dark at room temperature for 5 days. The seedlings are transplanted to soil in pot (4-6 inch) and maintained in greenhouse until blooming for sample collection. The temperature of greenhouse is kept around 27°C during daytime (12 hours) and 25°C for nighttime (12 hours). The daytime light is controlled to the level of 500 μmol m^-2^s^-1^. Rice plants are irrigated with deionized water every day and fertilized twice each week by filling the headspace in pots with 300-350 ppm Nitrogen (Jack’s 20-10-20).

### Isolation of rice egg cells and zygotes

The previous description was followed (Anderson et al 2013 and 2017, and Li et al 2019) with minor modifications. Six-ten mature and still closed florets (in which anther occupies most of the floret prior to anthesis) for egg cell isolation or pollinated open florets for a certain time for zygote isolation are collected into 0.4M mannitol in a 10 mL petri dish in the morning (9-11 am). The isolated cells were captured using a special micropipette and immediately used for fluorescent microscopy or temporally stored in a moist chamber for transfection with plasmid DNA.

### PCR cloning for cellular localization in rice cells

The mRNA from young rice flowers (Kitaake) are purified using Oligotex mRNA Mini Kit (Qiagen 70022) and cDNAs are synthesized with RevertAid First Strand cDNA Synthesis Kit (Thermo Scientific Inc, K1621). For the products of coding sequence (CDS) from the PCR reactions of rice *KRP5* (LOC_Os03g04490.1), *KRP4* (LOC_Os10g33310.1), and the F-box gene, *FB3* (LOC_Os08g09750), we used three pairs of specific primers (Supplementary Table S2) with Q5 High Fidelity DNA Polymerase (NEB, M0491) or Phusion High Fidelity DNA Polymerase (Thermo Scientific Inc F530S). Then, *FB3* CDS product is ligated to the vector pE3150 (Lee and Gelvin 2014; Lee et al 2008) at XhoI and HindIII to fuse with Enhanced Yellow Fluorescent Protein (EYFP) at C’ terminus, *KRP4* at EcoRI and SalI, *KRP5* at EcoRI and SalI. In addition, mCherry with nuclear localization signal peptides in the vector pE3275 was used as the positive control in the transfection for the nuclear localization.

### Isolation and transfection of rice leaf protoplasts

To test if the construct of the rice gene fused with EYFP work or not prior to the formal transfection of valuable isolated rice gametes and zygotes, the rice leaf protoplasts are prepared and transfected according to the procedure described by Wang et al (2013).

### Transfection of isolated rice egg cells and zygotes

The procedure was developed and optimized with reference to the previous (Koiso et al 2017 and Toda et al 2019). Briefly, we transferred the isolated cells (3 - 5, stored in the moist chamber) on to a micro slide (within a moisture chamber) in 8 µl MMG (4 mM Mes-KOH, pH5.7, 15 mM MgCl_2_ and 0.4 M mannitol), added with 2 µl plasmid DNA (1 µg, pE3275 and EYFP fused KRP or Fb3 construct) and 10 µl of 30% PEG with 100 mM CaCl_2_ in 0.4 M mannitol; the mixture was gently mixed twice using the micro-pipet and incubated at room temerature for 10 min, then the cells were carefully washed 3 times with fresh MMG (capillary replacement of ∼ 10 µl each time), followed by 3 more washes in modified W5 solution (154 mM NaCl, 125 mM CaCl_2_, 5 mM KCl, 2 mM MES; plus 500 mM Glucose). At last, we cultured the cells with 10 µl of the modified W5 solution containing Ampicillin (100 µg/ml) within the moist chamber at 25°C in dark for overnight up to 18 hours. For fluorescent microscopy, the droplet containg the cells on the slide was surrounded with cream (Petrolatum, Fisher Scientific, P66-1) as a square chamber and slowly covered with a coverslip.

### Fluorescent microscopy and image processing

Aniline blue fluorescence is used to monitor the pollen germination on stigma and growth along the style. For this purpose, the stigma and style assocaitd with the ovaryupper portion are stained in Aniline blue solution (0.005% in 0.15M K_2_HPO_4_ at pH8.2) for 10 min at room temperature., then viewed under the fluorescent microscope Axiovert 10 (Zeiss). To examine the viability and observe the nuclear, isolated gametes and zygotes or pollen grains are stained with Fluorescine Diacetate (FDA, excited at 48-510 nm and emited at 535-585 nm) and 4′, 6-Diamidino-2-phenylindole (DAPI, excited at 358 nm and emited at 461nm), separately, as follows: Made a square chamber on a clean slide with the Petrolatum cream and transfer isolated cells in 5-10 µl 0.4 M mannitol into it; Add the same volume of 2x FDA or DAPI staining solution which is diluted in 1:1000 from the stock (1µg/µl) in 0.4M Mannitol; Keep the slide in a moist chamber (a petri-dish with a wet paper tower) in dark for 3-5 minutes for FDA, or ∼ 30 minutes for DAPI; Put a cover slip over the cream chamber for microscopy (Axiovert 10). To observe transfected rice cells, Nikon Eclipse Ni matched with the light source of X-Cite 120LED is used with appropriate filters for EYFP (excitation at 514 nm and emission at 527 nm) and for mCherry (excitation at 587 nm, emission at 610 nm). The images were processed with ImageJ.

## Results and discussion

### Isolation of rice egg cells and zygotes

To isolate rice zygotes, we adopted the same protocol for isolating rice egg cells, but did polination prior to the floret collection at the desired time. For self polination, the selected mature florets are carefully open with the aid of tweezers. In a couple of minutes, we start timing and slightly flicked those open florets to help pollen grains fall onto the stigma. For cross pollination, we follow the procedure described by Dr. Susan McCouch, Department of Plant Breeding and Genetics, Cornell University (http://ricelab.plbr.cornell.edu/cross_pollinating_rice), including anther removal from the recipient florets (emasculation) and 3-5 times of quick addition of the pollen freshly collected from the donor plants.

The representative images of isolated rice zygotes from three different stages (3, 6 and 9 HAP) are shown as in Fig. 1. They have the similar morphology to the isolated egg cells except slightly smaller size. As demonstrated in FDA staining, they are viable for hours after isolation if cultured in the appropriate solution or culture medium. Like the isolated egg cells the isolated zygots have much larger nuclear with active euchromatin.

**Fig. 1.**
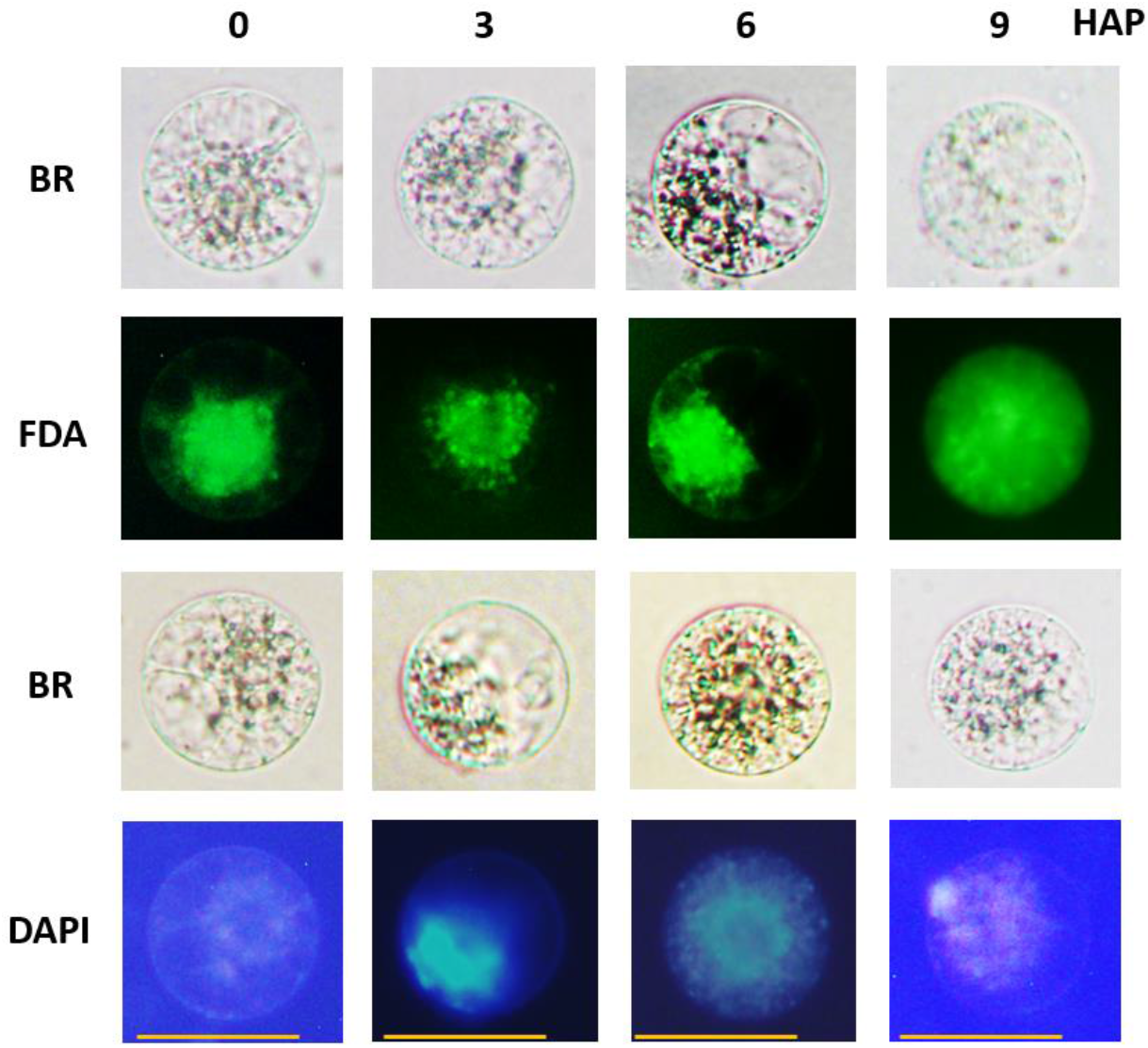
Isolated rice egg cells and zygotes stained with FDA and DAPI. Representative images of isolated rice egg cells (0 HAP) and zygotes at different developmental stages (3, 6 and 9 HAP). BR, differential interference contrast images of isolated cells; FDA (Fluorescein diacetate) indicates the viability of isolated cells in green fluorescence; DAPI indicates nuclei of isolated cells in blue fluorescence. The observation had 3 biological replicates (N = 3-6 cells in each).

One of common concerns of the isolated zygotes is, if any, how many unfertilized egg cells could be mixed with the fertilized zygotes in the isolation. Since the marker gene method does not help address this issue, we adopted two other different measures. One is the microscopy with Aniline Blue fluorescence to isolate the self zygotes at 2 HAP (Supplemental Fig. S1). Once finding the pollen tube(s) grwing along the stigma which indicates the higher chance of fertilization, the corresponding lower portion ovary will be taken for zygote isolation. The other is seed setting assay (Supplemental Table S1). Under the condition we set in greehouse for rice growth, the seed set rate is 97%, which told us that, if any, the rate of unfertilized egg cells is lower than 3% in our isolated zygote sample; in contrast, the seed set rate is only 1% in the minus control of emasqulated florets without pollination. In conclusion, the quality of our isolated zygotes from self-pollination is reliable for the transfection for cellular localization and Bimolecular Fluorescent Complementation (BiFC), as it was previously used for the studies of rice gametic and zygotic siRNA (Li et al. 2020 and 2022).

### Transient transfection of isolated rice leaf protoplasts with EYFP-fused rice cell cycle factors

According to the rice genomic database data, three rice genes, *KRP4, KRP5* and *FB3*, encode putative cell cycle factors, which are likely important players in the control of rice zygotic cell division, the formal initiation of rice embryogenesis. Therefore, we are interested in testing their subcellular localization in isolated rice egg cells and zygotes. To ensure that the fluorescent fused constructs and the transfection procedure work prior to using the valuable isolated rice cells, we took rice leaf protoplasts as the pretrial material since they are widely used before in analysis of protein-protein interactions (PPI) and cellular localization (Lv Q et al. 2014; Shi et al. 2019; Wang et al. 2013).

As shown in Figs. 2a and b, the typical isolated rice leaf protoplast presents as transparent sphere in size of 30 ∼ 50 µm. Due to the pressure of big central vacuoles, their nuclei are pushed to plasma membrane region (the edge of 2D image) and accurately localized by transfection with mCherry_NSL_ (red signal). In the minus control (Fig. 2a, the top image #3 from left), EYFP is expressed in both nuclear and protoplasm, but both EYFP-KRP5 and EYFP-KRP4 are only expressed in nuclei (Fig. 2a) and this rate is up to over 90% (Fig. 2c), indicating KRP5 and KRP4 are nuclear proteins, which are consistent with their putative molecular function in the cell cycle control. We also observed that EYFP-Fb3 is expressed in either nuclei or cytoplasm with about 50% of chance for each (Figs. 2b and c), and each truncated Fb3 fused with EYFP has higher chance of expression in cytoplasm, indicating both F-box domain and LRR region contribute to the determination of FB3 localization in rice protoplasts.

**Fig. 2.**
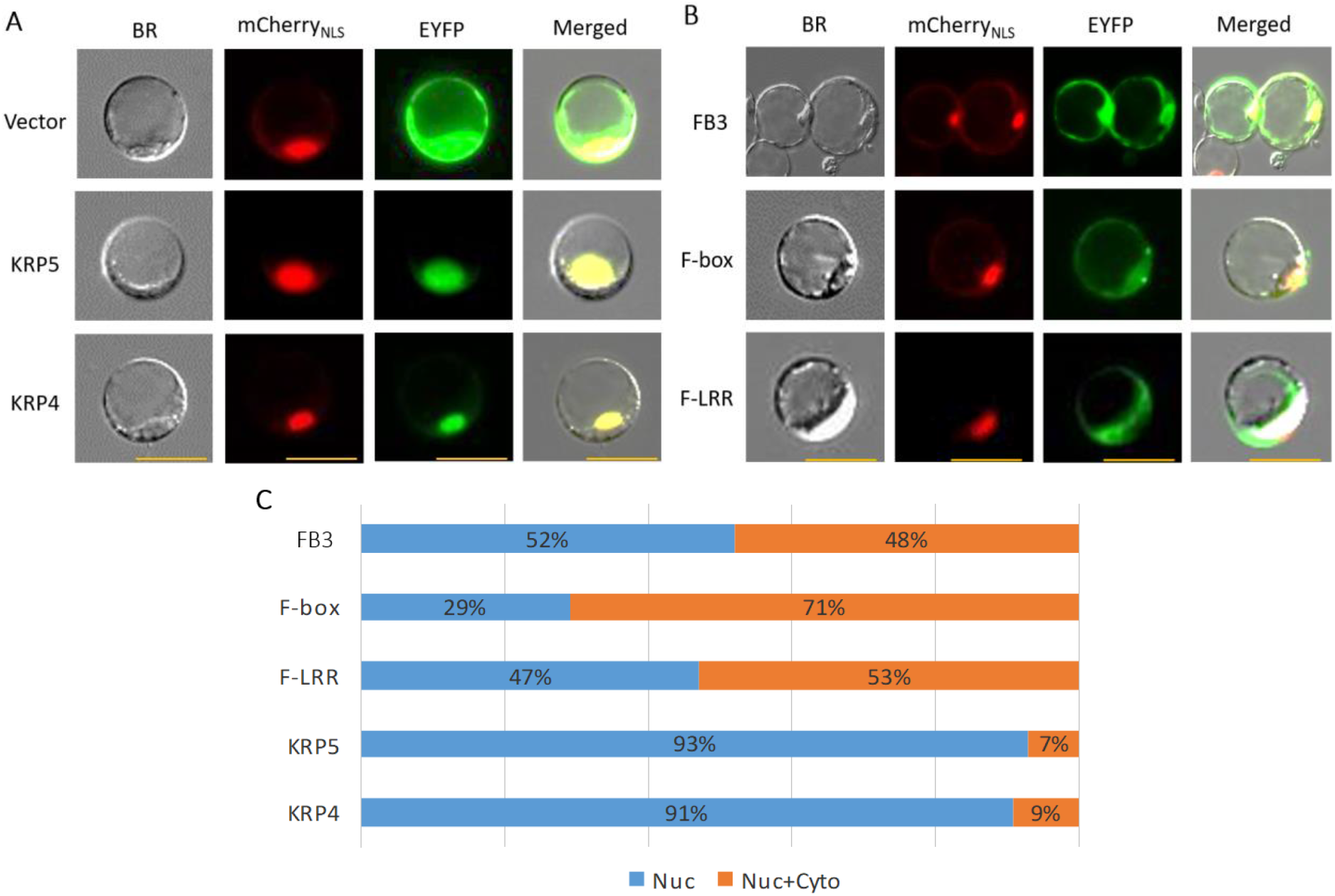
Cellular localization of protein KRP5, KRP4 and FB3 in rice leaf protoplasts. **a** Protoplasts were transfected with *KRP5* and *KRP4* gene fused with *EYFP* at 5’-terminus (*EYFP-KRP5* and *EYFP-KRP4*) driven by cauliflower mosaic virus (CaMC) 35S promoter; Vector (carrying *EYFP*) as the control. BR represents images of bright field. The mCherryNLS represents mCherry linked with Nuclear Localization Sequence (NLS, 12 amino acid residues) and was used as the nuclear marker. EYFP shows the cellular localization of protein KRP5 and KRP4. Merged refers the overlaid image from the other three in each transfection. Scale bars: 50 µm. **b** Protoplasts were transfected with *EYFP* fused *FB3, F-box* (300bp from 3’ terminus of *FB3* with F-box domain) and *F-LRR* (1.4 kb of *FB3* from 5’ terminus with Leucine Rich Region) at N-terminus. Scale bars: 50 µm. **c** Localization frequency of EYFP fused rice genes (*KRP4, KRP5* and *FB3*) and *FB3* fragments (*F-box* and *F-LRR*). Blue bars represent the percentage of protoplasts with Nuclear (Nuc) localization; orange bars indicate the percentage of protoplasts with nucleus-cytoplasm (Nuc+Cyto) localization.

In a word, the constructs of EYFP-KRP5, EYFP-KRP4 and EYFP-Fb3 as well as the procedure for transient trasfection work well for the cellular localization in rice protoplasts, and hopefully they will also work in isolated rice egg cells and zygotes.

### Transfection of isolated rice egg cells and zygotes with EYFP-fused rice cell cycle factors

Although the above constructs and procedure work well in transfecting rice protoplasts, it is difficult to apply them to isolated rice egg cells and zygotes. These isolated cells are not only limited in number, and also very vulnerable to multiple transferring and washing during the process of transfection. Therefore, the special caution was taken to carry out the procedure stated in method part.

As a result, the transfection worked through in the isolated rice egg cells and zygotes. As shown in Fig. 3a and b, mCherry_NLS_ expressed in all nuclei (red signal) of the egg cells (0 HAP) and zygotes at 2 HAP and 9 HAP; the co-transformed EYFP-KRP5 and EYFP-KRP4 also expressed in the similar pattern (green signal). The merged images demonstrate the nuclear localization of KRP5 and KRP4 in rice egg cells and zygotes, consistent with our observation in the transfected rice protoplasts. However, as shown in Fig. 3c, the expression patterns of mCherry_NLS_ (red signal) and EYFP-Fb3 (green signal) in the transfected cells are not overlapped well, indicating the localization of EYFP-Fb3 mainly in nuclei and partially in cytoplasm.

**Fig. 3.**
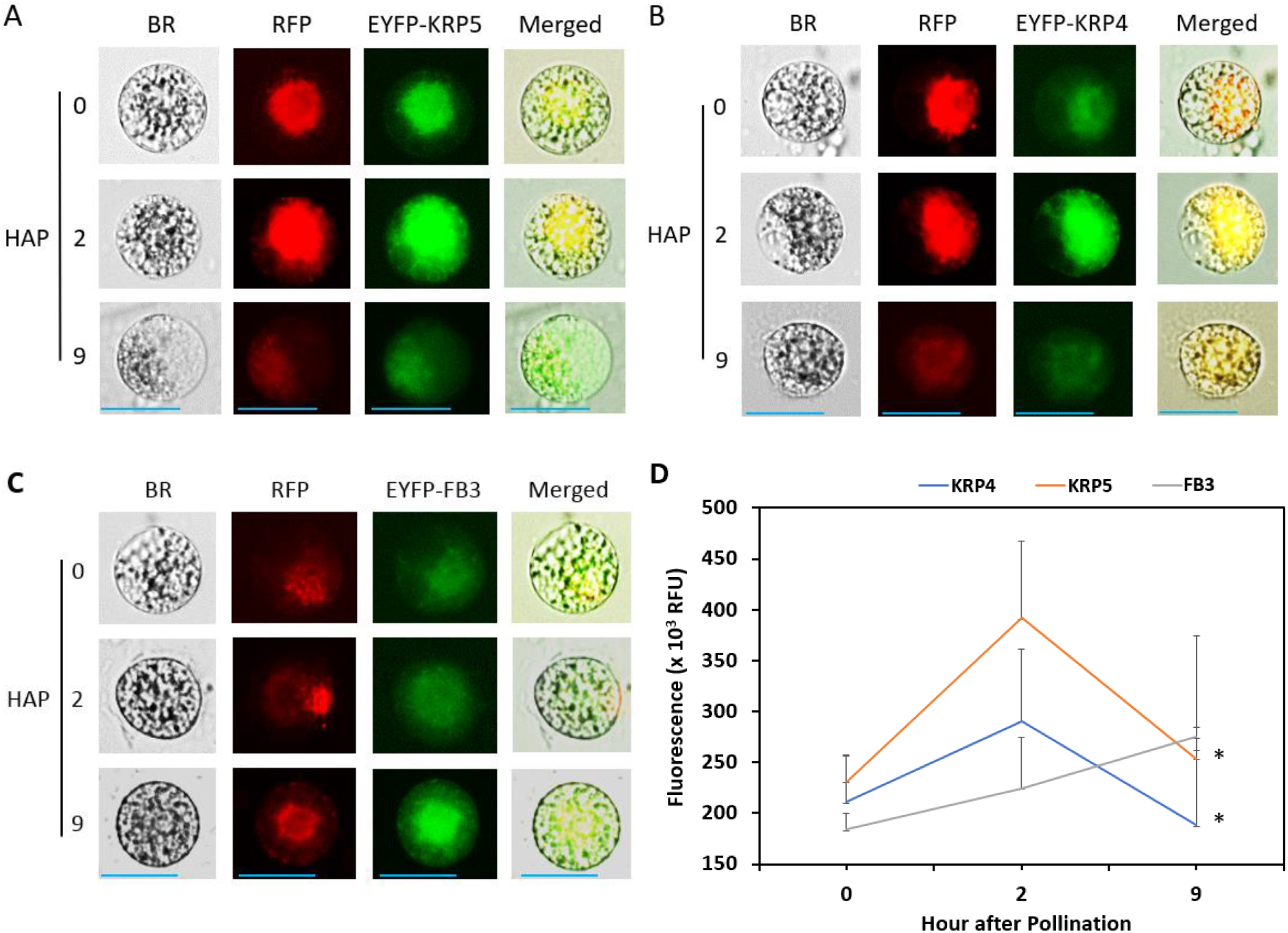
Transfection of isolated rice egg cells and zygotes for cellular localization with EYFP-fused 3 putative rice cell cycle genes. **a** Representative images of rice egg cells and zygotes transfected with *EYFP-KRP5*. **b** Representative images of rice egg cells and zygotes transfected with *EYFP-KRP5*. **c** Representative images of rice egg cell and zygotes transfected with *EYFP-FB3*. In a-c, HAP, hour after pollination. BR, cell images in gray under bright field. RFP represents mCherry linked with Nuclear Localization Sequence (NLS) as the nuclear marker. Merged, overlapping all other images in the same transfection. Scale bars in a-c: 50 µm. **d** Protein expression profiles of KRP4, KRP5 and FB3 in rice egg cells and zygotes. Fluorescent intensity was quantified with ImageJ. Values are the mean and error bars are the standard deviation for three biological replicates (N = 3-6 cells in each). The * indicates a significant difference in fluorescence of KRP4 or KRP5 between 2 HAP and 9 HAP via one-way ANOVA analysis (*P* < 0.05).

It is also interesting to compare the fluorescent intensity of the transfected cells at different developmental stages. In Fig. 3a and b, both EYFP-KRP5 and EYFP-KRP4 give stronger signal in the egg cells and zygotes at 2 HAP (both in G1 phase of cell cycle) but weaker in zygotes of 9 HAP (S-G2 phase of cell cycle). In contrast to Fig. 3c, the expression level of EYFP-FB3 in the egg cell and zygote of 2 HAP (the G1 phase) is obviously lower than that of zygote at 9 HAP (the G2 phase). This feature is confirmed as shown in Fig. 3d. It indicates the putative function of these proteins in plant zygotic cell cycle control, and demonstrates this transient transfection system as a potentially useful approach in testing the specific protein expression level in plant gametes and zygotes.

The same procedure for cellular localization also has been applied in Bimolecular Fluorescent Complementation (BiFC) in isolated rice egg cells and gametes at different developmental stages and it is successful (Xu and Russell 2021). Put these two parts of assays together, we totally used 546 isolated egg cells and zygotes, among which the first 128 was consumed in the pre-test and the rest 418 for the formal transfection. From the latter, 148 transfected cells survived through the overnight incubation for protein expression and 86 of these cells were observed the expected fluorescent signals. Thus, the surviving rate after transfection is 35.4% and successful co-transfection rate is 20.5%, an efficiency not lower than what we expected. To our knowledge, this is the first trial of the transient transfection to observe important protein expression using hundreds of isolated living rice egg cells and zygotes.

### Advantageous features of isolated rice gametes and zygotes for the study in plant biology

From this work we recognized four advantageous features of isolated rice egg cells and zygotes to serve as an ideal system for research in plant biology.

#### 1. Transparency

Plant zygotes and gametes contain plastids but chloroplast-free and those single cells isolated under our condition are at least partially cell wall free. They are suitable for fluorescent microscopy to detect labeled gene expression without auto fluorescent interference.

#### 2. Purity

The isolated rice gametes and zygotes are very pure, as verified by PCR using marker genes (Li et al. 2019) and the study of siRNAs (Li et al. 2020 and 2022). This feature makes it possible to study the epigenetic modifications in rice gametes and zygotes (Khanday and Sundaresan 2021) to reveal the mechanism underlying the zygotic development.

#### 3. Viability

As shown in cell images of FDA staining (Fig. 1), all isolated rice egg cells and zygotes are viable. In our cellular localization, over 35% of transfected rice egg cells and zygotes are living after overnight incubation. That is why the isolated rice egg cells and zygotes have been used for PEG-Ca^++^ mediated transfection and tissue culture for permanently transformed rice plants (Koiso et al. 2017; Toda et al. 2019). Using this technique, we may get specific cell marker lines like those in Arabidopsis (Chamberlin and Lawi 2017) to facilitate the study of rice seed biology.

#### 4. Integrity

Since no cell-wall digestion enzyme is used in our manipulation, the isolated rice egg cells and zygotes are intact and advantageous for studying surface glycoproteins that may be involved in the fertilization-recognition or zygotic development in crop plants. Studies in mammals have revealed that, the sugar chains of glycoproteins play roles in cell-to-cell recognition and fusion (Avella et al. 2013; Xu and Tsao 1997). During the past decades, several gamete specific proteins (e.g. GEX2, HAP2/GCS1, DMP8/9, EC1 and ECS) have been functionally characterized for their involvement in the cell-to-cell recognition during the double fertilization in Arabidopsis (Cyprys et al. 2019; Dresselhaus et al. 2016; Mori et al. 2014; Sprunck 2020; Yu et al. 2021), but it is unknown if they are glycoproteins and if they are universal in the double fertilization of crop plants.

Therefore, our approach to manipulating rice gametes and zygotes is important, unique, and informative for biological research.

## Author contribution statement

H.X. designed and performed the experiment, analyzed data, and drafted the manuscript. L.B. assisted with PCR performance and participated in manuscript preparation. M.L. provided vectors and participated in molecular cloning and manuscript preparation. V.S. coordinated and supported the project and provided instructive suggestions. H.F. managed rice plant growth and participated in the cell isolation. S.R. supervised and supported the research, managed rice cell isolation, and participated in manuscript preparation.

## Acknowledgements

We thank Mr. Joshua Chesnut and Dr. Daniel Jones for demonstrating manual isolation of rice egg cells, and Dr. Ben Smith and Dr. David Thomas for instructions in epi-fluorescence microscopy. The authors thank the National Science Foundation (USA) for financial support (Grant #IOS-1547760, to Dr. Venkatesan Sundaresan and Dr. Scott Russell).

