## Supplemental Information for "Isolation and Transfection of Rice Egg Cells and Zygotes for Cellular Localization"

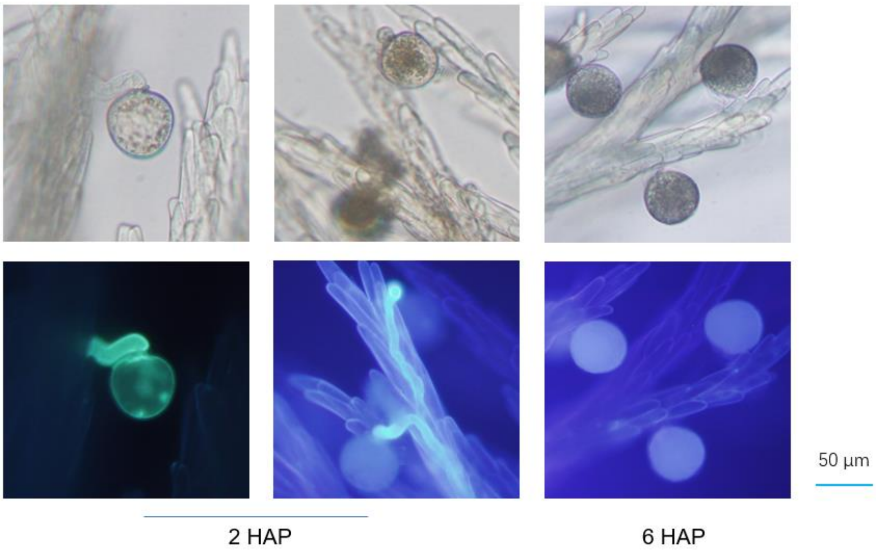


Fig. S1 Rice pollen grain and pollen tube located on stigma and grown along the style. The upper portion of carpel is stained with Aniline Blue and observed under fluorescent microscope. HAP: hour after pollination.

**Table S1** Rice seed-set assay control

| | Emasculated without pollination | | | Self-pollinated | | |
|---|---|---|---|---|---|---|
| Index# | Emasculated florets | Seeds | Seed-set(%) | Pollinated florets | Seeds | Seed-set (%) |
| 1 | 11 | 1 | | 9 | 8 | |
| 2 | 20 | | | 7 | 7 | |
| 3 | 21 | 1 | | 9 | 9 | |
| 4 | 24 | | | 8 | 8 | |
| 5 | 22 | | | 10 | 9 | |
| 6 | 13 | | | 12 | 12 | |
| 7 | 14 | | | 11 | 10 | |
| 8 | 12 | | | 8 | 8 | |
| 9 | | | | 9 | 9 | |
| 10 | | | | 10 | 10 | |
| 11 | | | | 8 | 8 | |
| Total | 137 | 2 | 1 | 101 | 98 | 97 |

**Table S2** Primers for cellular localization constructs

| Primers | Sequences 5'-3' | Product size (bp) |
|---|---|---|
| FB3-EYFP-F-XhoI | TGTCTCGAGCTATGCTCGGGAGGAACGCCAT | |
| FB3-EYFP-R-HindIII | TGCAAGCTTCGGAACTCCATATCTGAATCCTC | 1684 |
| KRP4-EYFP-F-EcoRI | GTCGAATTCTATGGGCAAGTACATGCGCAA | |
| KRP4-EYFP-R-SalI | TGCGTCGACGTCTAGCTTGACCCATTCAAA | 609 |
| KRP5-EYFP-F-EcoRI | GTCGAATTCTATGGGGAAGTACATGCGGAAG | |
| KRP5-EYFP-R-SalI | TGCGTCGACGCAGTCTAGCCTTGTCCATTC | 690 |
